# Analyzing cancer gene expression data through the lens of normal tissue-specificity

**DOI:** 10.1101/2021.01.25.428166

**Authors:** H. Robert Frost

**Affiliations:** Department of Biomedical Data Science, Geisel School of Medicine, Dartmouth College, Hanover, NH 03755

## Abstract

The genetic alterations that underlie cancer development are highly tissue-specific with the majority of driving alterations occurring in only a few cancer types and with alterations common to multiple cancer types often showing a tissue-specific functional impact. This tissue-specificity means that the biology of normal tissues carries important information regarding the pathophysiology of the associated cancers, information that can be leveraged to improve the power and accuracy of cancer genomic analyses. Research exploring the use of normal tissue data for the analysis of cancer genomics has primarily focused on the paired analysis of tumor and adjacent normal samples. Efforts to leverage the general characteristics of normal tissue for cancer analysis has received less attention with most investigations focusing on understanding the tissue-specific factors that lead to individual genomic alterations or dysregulated pathways within a single cancer type. To address this gap and support scenarios where adjacent normal tissue samples are not available, we explored the genome-wide association between the transcriptomes of 21 solid human cancers and their associated normal tissues as profiled in healthy individuals. While the average gene expression profiles of normal and cancerous tissue may appear distinct, with normal tissues more similar to other normal tissues than to the associated cancer types, when transformed into relative expression values, i.e., the ratio of expression in one tissue or cancer relative to the mean in other tissues or cancers, the close association between gene activity in normal tissues and related cancers is revealed. As we demonstrate through an analysis of tumor data from The Cancer Genome Atlas and normal tissue data from the Human Protein Atlas, this association between tissue-specific and cancer-specific expression values can be leveraged to improve the prognostic modeling of cancer, the comparative analysis of different cancer types, and the analysis of cancer and normal tissue pairs.

## 1 Background

The biology of human cancer is highly tissue and cell type-specific [1–4]. Most cancer driver genes are altered in only a small number of cancer types and, for drivers that are broadly mutated, the impact of the alteration often varies significantly between tissue types. Examples of genes with a tissue-specific pattern of alteration include von Hippel Lindau tumor suppressor (VHL) in renal cancer, adenomatous polyposis coli (APC) in colorectal cancer, and KRAS in pancreatic, lung and colorectal cancers. Examples of driver genes that are altered in multiple cancer types but have a tissue-specific functional impact include BRAC1/BRAC2, which are impacted by germ-line mutation but lead to cancer primarily in estrogen-sensitive tissues (e.g., breast and ovaries), and BRAF, which can be effectively inhibited in BRAF-mutated melanoma but not in BRAF-mutated colon cancer. Only a small number of driver genes are broadly altered in many different cancers with similar functional consequences, e.g., TP53 and MYC. The tissue specificity of cancer driver genes is due to both cell intrinsic factors (i.e., epigenetic landscape and corresponding regulatory circuitry) and cell extrinsic factors (cell-cell interactions in the tissue microenvironment and environmental exposures). One important cell intrinsic factor is the cell type-specificity of proliferation drivers; 80-90% of the genes that drive proliferation function in only a limited number of cell types and these genes are frequently impacted by aneuploidy in cancer [2,3]. Examples of cell extrinsic factors include exposure to estrogen in breast and ovarian tissue and the consequent vulnerability to BRAC1/BRAC2 mutations, and exposure to UV radiation in melanoma which leads to both increased sensitivity to the alteration of nucleotide excision repair genes and to an increase in the number of neoantigens and improved response to immunotherapy [4].

Given the significant tissue-specificity of cancer, the pattern of gene activity within normal tissue carries important information about the developmental trajectory of associated cancers and how those tumors will respond to therapeutic interventions. Efforts to leverage this information for the analysis of cancer genomics data have primarily focused on the analysis of paired tumor and adjacent normal tissue samples from repositories such as The Cancer Genome Atlas (TCGA) [5]. Analyses of paired tumor/normal data includes the work by Huang et al. exploring cancer prognosis prediction using data from adjacent normal samples [6] and research by Hu et al. investigating the pattern of relative gene expression between tumors and adjacent normal tissue [7]. Although the use of paired normal tissue data can improve the analysis of tumor data, this approach has two important limitations. First, adjacent normal tissue is impacted by the tumor microenvironment so is not accurate reflection of the biology of that tissue in a healthy individual. This so-called field cancerization effect explains why genomic data from adjacent normal tissue can independently predict cancer survival and, in some cases, provide superior performance relative to models based on tumor-derived data [6]. The second limitation is the fact that data from adjacent normal tissue samples is often not available.

The limitations of paired tumor/normal data analysis motivate the exploration of associations between the general features of normal tissues, as measured in healthy individuals, and the characteristics of the cancer types that can develop in those tissues. While many researchers have investigated the association between normal tissue biology and cancer development in the context of specific cancer types and the associated cancer drivers (e.g., the association between estrogen sensitive tissues, BRAC1/BRAC2 mutations and cancer development), only a limited number of researchers have explored the genome-wide association between normal tissue and cancer gene activity across multiple cancer types. The most prominent recent investigation into the general relationship between normal tissue and cancer gene activity is the work by the Uhlen et al. [8], who analyzed the association between gene expression in human solid tumors profiled by the TCGA and the corresponding normal tissues profiled by the Human Protein Atlas (HPA) [9]. Although the work by Uhlen et al. evaluated a large number of tumor types and normal tissues, the analysis focused on the association between mean gene expression in tumors and mean expression in the corresponding normal tissue, which revealed that the transcriptomes of normal tissues and cancers form two distinct clusters with each normal tissue more similar to other normal tissues than to the corresponding cancer type (see Uhlen et al. Figure S1). Successful use of normal tissue data for cancer genomics analysis was limited to liver cancer, where it was found that genes with elevated expression in normal liver tissue are favorably prognostic in liver cancer and genes whose expression was not specific to liver tissue are unfavorably prognostic. A similar liver-specific analysis was conducted by Li et al. [10], which revealed a concordant finding regarding the prognostic power of liver-specific genes.

To address the gap in effective pan-cancer approaches for leveraging normal tissue gene activity for the analysis of cancer genomics data, we explored the genome-wide association between the transcriptomes of 21 solid human cancers and their associated normal tissues as profiled in healthy individuals. While the average gene expression profiles of normal and cancerous tissue may appear distinct, as found by Uhlen et al. [8] with normal tissues more similar to other normal tissues than to the associated cancer types, when transformed into relative expression values, i.e., the ratio of expression in one tissue or cancer relative to the mean in other tissues or cancers, the close association between gene expression in normal tissues and related cancers is revealed. As we demonstrate through an analysis of tumor data from TCGA and normal tissue data from the HPA, this association between tissue-specific and cancer-specific expression values can be leveraged to improve the prognostic modeling of cancer, the comparative analysis of different cancer types, and the analysis of cancer and normal tissue pairs.

## 2 Results

### 2.1 Analysis overview

The results presented in this paper are based on bulk RNA-seq data from TCGA for 21 human solid cancers and from the HPA for the associated 18 normal human tissues (see Table 2 below for a list of the cancer types and matching normal tissues). These gene expression data sets were used to compute several statistics that capture different aspects of gene activity and prognostic value. So that readers can more easily follow the main results presented in Sections 2.2 - 2.7, a concise definition of these statistics is included in Table 1.

**Table 1:** Gene-level statistics computed on TCGA and HPA RNA-seq gene expression data and TCGA survival data.

| Statistic | Description |
| --- | --- |
| $c_{i,j}$ | Mean expression of gene $i$ in cancer type $j$. The average of these mean expression values for gene $i$ across all 21 cancer types is represented by $\bar{c}_i$. |
| $n_{i,j}$ | Mean expression of gene $i$ in normal tissue $j$ (the tissue associated with cancer type $j$). The average of these mean expression values for gene $i$ across all 18 normal tissues is represented by $\bar{n}_i$. |
| $c_{i,j}^* = \log_2(c_{i,j}/\bar{c}_i)$ | Cancer-specific expression of gene $i$ in cancer type $j$ as computed by the log fold-change in expression of gene $i$ between cancer type $j$ and the average in all cancer types. |
| $n_{i,j}^* = \log_2(n_{i,j}/\bar{n}_i)$ | Normal tissue-specific expression of gene $i$ in tissue $j$ as computed by the log fold-change in expression of gene $i$ between tissue $j$ and the average in all normal tissues. |
| $r_{i,j} = \log_2(c_{i,j}/n_{i,j})$ | Relative expression of gene $i$ between cancer type $j$ and the corresponding normal tissue as computed by the log fold-change in mean expression of gene $i$ in cancer $j$ and normal tissue $j$. |
| $s_{i,j}$ | Cancer prognostic value of gene $i$ for cancer type $j$ as computed by the signed log of a p-value generated by Uhlen et al. [8] to capture the significance of the association between expression of gene $i$ and survival for cancer type $j$. For genes with a favorable prognostic value, the -log of the p-value is used to produce positive values. For genes with an unfavorable prognostic value, the log of the p-value is used instead to generate negative values. |

**Table 2:** The 21 analyzed TCGA cancer types and corresponding HPA normal tissues. The ‘Cancer/normal correlation’ column contains the Spearman rank correlation between the mean gene expression values in the TCGA cancer samples (i.e., the *c_i,j_* statistics) and the mean expression values in the associated HPA normal tissue (i.e., the *n_i.j_* statistics). The ‘Most correlated’ column lists the HPA normal tissue whose mean expression had the highest rank correlation with cancer mean expression. A value of ‘*’ indicates that the associated normal tissue had the largest correlation.

| TCGA abbrev. | Cancer type | HPA tissue | Cancer/normal correlation | Most correlated |
| --- | --- | --- | --- | --- |
| BLCA | Bladder Urothelial Carcinoma | urinary bladder | 0.901 | * |
| BRCA | Breast Invasive Carcinoma | breast | 0.924 | * |
| CESC | Cervical Squamous Cell Carcinoma and ... | cervix, uterine | 0.852 | urinary bladder |
| COAD | Colon Adenocarcinoma | colon | 0.93 | * |
| GBM | Glioblastoma Multiforme | cerebral cortex | 0.895 | * |
| HNSC | Head and Neck Squamous Cell Carcinoma | tonsil | 0.862 | skin |
| KICH | Kidney Chromophobe | kidney | 0.932 | * |
| KIRC | Kidney Renal Clear Cell Carcinoma | kidney | 0.931 | * |
| KIRP | Kidney Renal Papillary Cell Carcinoma | kidney | 0.925 | * |
| LIHC | Liver Hepatocellular Carcinoma | liver | 0.929 | * |
| LUAD | Lung Adenocarcinoma | lung | 0.925 | * |
| LUSC | Lung Squamous Cell Carcinoma | lung | 0.886 | urinary bladder |
| OV | Ovarian Serous Cystadenocarcinoma | ovary | 0.817 | cervix, uterine |
| PAAD | Pancreatic Adenocarcinoma | pancreas | 0.886 | stomach |
| PRAD | Prostate Adenocarcinoma | prostate | 0.951 | * |
| READ | Rectum Adenocarcinoma | rectum | 0.912 | colon |
| SKCM | Skin Cutaneous Melanoma | skin | 0.838 | urinary bladder |
| STAD | Stomach Adenocarcinoma | stomach | 0.921 | * |
| TGCT | Testicular Germ Cell Tumors | testis | 0.698 | urinary bladder |
| THCA | Thyroid Carcinoma | thyroid gland | 0.934 | * |
| UCEC | Uterine Corpus Endometrial Carcinoma | endometrium | 0.868 | cervix, uterine |

### 2.2 Association between gene activity in normal and neoplastic tissue

Figure 1 illustrates the projection of the 21 analyzed cancer types and 18 corresponding normal tissue types onto the first two principal components (PCs) from a principal component analysis (PCA) of mean expression values, i.e., PCA of a matrix that contains all *c_i,j_* and *n_i.j_* statistics. As seen in Figure 1 and in Figure S1 from the Ulhen et al. [8] paper, cancers and normal tissues cluster separately according to gene expression data (i.e., cancers are more similar to each other than to the corresponding normal tissues). Liver cancer and normal liver tissue are noticeable outliers in this projection and, unlike most other profiled cancer/normal pairs, have mean transcriptomes that are more similar to each other than to other cancers or normal tissues. This strong association between normal liver and liver cancer gene expression was identified and explored by both Uhlen et al. [8] and Li et al. [10]. Despite the separation between cancer and normal tissue transcriptomes in the PCA projection, a very high correlation still exists between the mean gene expression values in a specific cancer and the expression values in the corresponding normal tissue. This association is captured in Table 2, which lists the rank correlation between cancer and associated normal tissue mean expression values, i.e., the *c_i,j_* and *n_i.j_* statistics. As shown in Table 2, the majority of the profiled cancers are most strongly correlated with their corresponding normal tissue.

**Figure 1:**
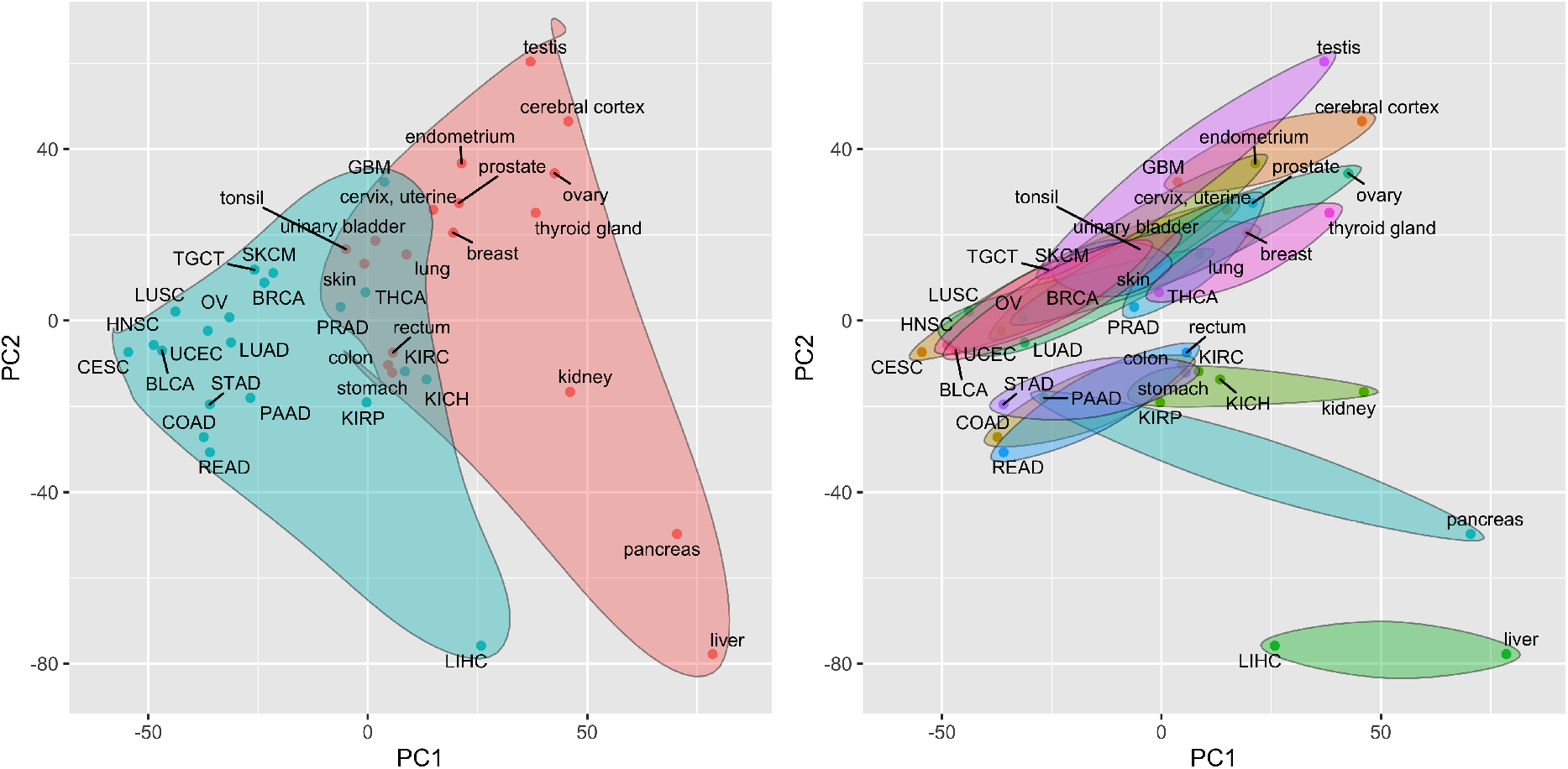
Projection of TCGA cancer types and associated HPA normal tissues onto the first two principal components. Principal components are computed from a matrix of mean gene expression values, i.e., a matrix containing all *c_i,j_* and *n_i.j_* statistics. a) Cancers are represented by blue points and are enclosed in the blue shaded region; normal tissues are represented by red points and are enclosed in the red shaded region. b) Each normal tissue and the associated cancer type(s) are enclosed in a separate shaded region.

### 2.3 Association between normal tissue-specificity and cancer type-specificity

When analyzing tissue-specific or cancer-specific gene expression, i.e., the ratio of expression in one tissue or cancer to the mean expression in all tissues or cancers as quantified by the 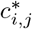 and 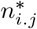 statis-tics, cancers and normal tissues no longer cluster separately and most cancers look very similar to the corresponding normal tissue. Figure 2 illustrates the projection of the analyzed cancers and normal tissues onto the first two PCs of the relative mean expression values, i.e., the PCs of a matrix combining all 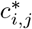 and 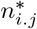 statistics. As seen in Figure 2, cancers and normal tissues no longer separate and each cancer tends to be very close to the corresponding normal tissue. These findings are quantified in Table 3 using the Euclidean distances between cancer and normal tissues in the space of the first two PCs. Specifically, the ‘Cancer/normal relative distance’ column contains the ratio of the distance between a given cancer and normal tissue in the PC space visualized by Figure 1 relative to average distance between that cancer and all other cancers or normal tissues. A relative distance of 1.0 indicates that the cancer mean expression values are no more similar to the expression values in the corresponding normal tissue than to a randomly selected cancer or normal tissue. Values above 1.0 indicate less similarity than would be expected at random and values below 1.0 indicate that the cancer and normal tissue transcriptomes are more similar than would be expected at random. The ‘Cancer/normal-specific relative distance’ column contains equivalent relative distance values generated in the space of the first two PCs computed on the cancer and tissue-specific expression data as visualized in Figure 2. The ‘Distance ratio’ column contains the ratio of the cancer/tissue-specific relative distances to the non-specific relative distances. Importantly, the distance ratios for most cancer/normal pairs are significantly below 1.0, i.e., cancer and normal tissue transcriptomic profiles are much more similar when evaluated in terms of cancer/tissue-specific values. Pancreatic cancer and skin cancer are two notable exceptions. For skin cancer, this result is likely due to the fact that the TCGA expression data is generated on metastatic lesions rather than primary tumors,which will reflect the properties of the host tissue. The implication of these results is that differences in gene expression between cancers largely reflect differences found between the corresponding normal tissues.

**Table 3:**
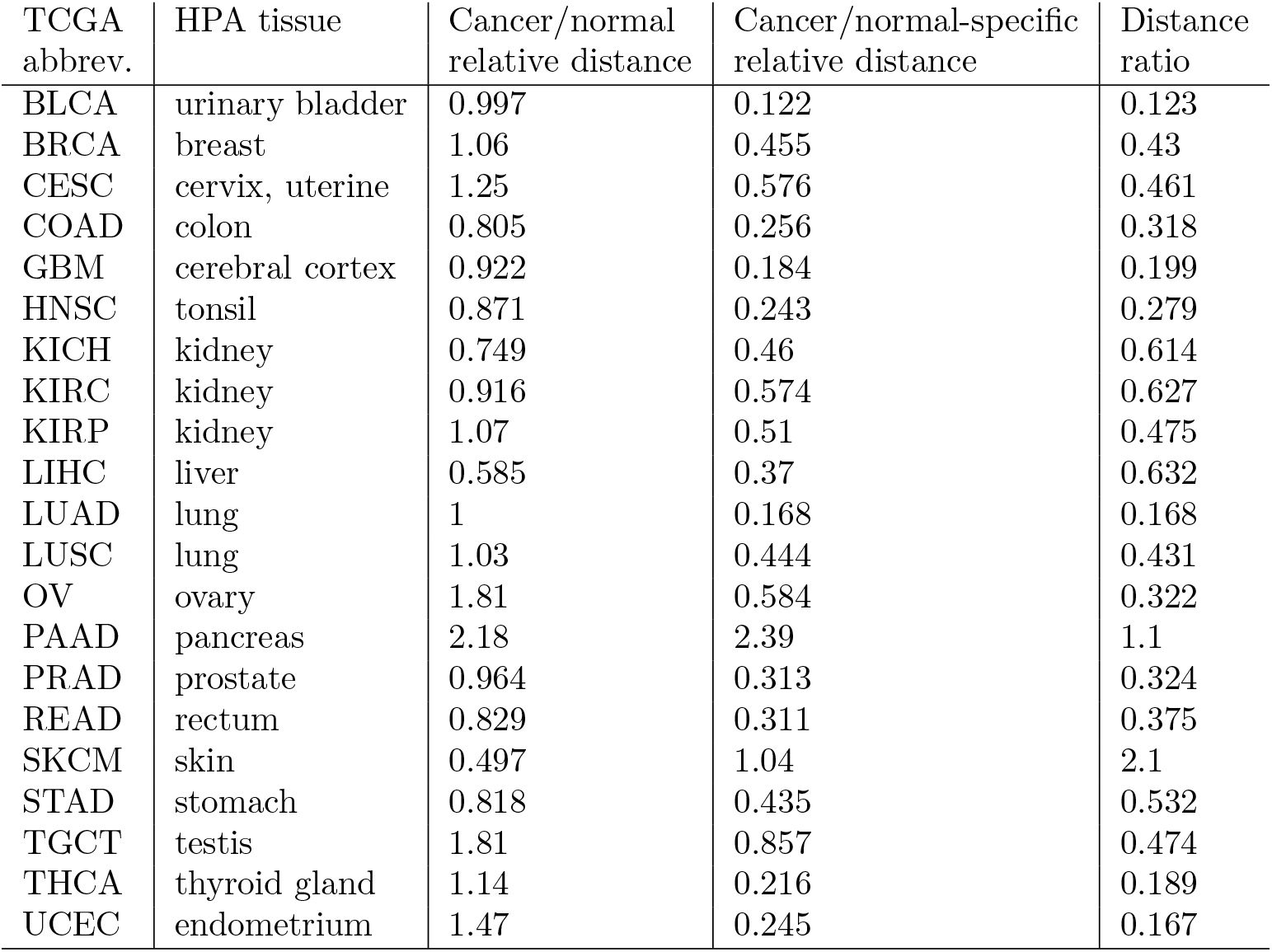
Distances between each cancer type and the corresponding normal tissue in reduced principal component (PC) space. The Euclidean distance was computed between the projections of each cancer type and the associated normal tissue in the space spanned by the first two PCs of the mean gene expression matrix (as shown in Figures 1 and 2). The ‘Cancer/normal relative distance’ column contains the ratio of the distance between each cancer and normal tissue pair as visualized in Figure 1 to the average distance between the cancer and all other cancers or normal tissues. The ‘Cancer/normal-specific relative distance’ column contains a similar distance ratio computed in the space spanned by the first two PCs of cancer-specific and normal tissue-specific mean expression values as visualized in Figure 2. The ‘Distance ratio’ column contains the ratio of the ‘Cancer/normal-specific relative distance’ to the ‘Cancer/normal relative distance’.

| TCGA abbrev. | HPA tissue | Cancer/normal relative distance | Cancer/normal-specific relative distance | Distance ratio |
| --- | --- | --- | --- | --- |
| BLCA | urinary bladder | 0.997 | 0.122 | 0.123 |
| BRCA | breast | 1.06 | 0.455 | 0.43 |
| CESC | cervix, uterine | 1.25 | 0.576 | 0.461 |
| COAD | colon | 0.805 | 0.256 | 0.318 |
| GBM | cerebral cortex | 0.922 | 0.184 | 0.199 |
| HNSC | tonsil | 0.871 | 0.243 | 0.279 |
| KICH | kidney | 0.749 | 0.46 | 0.614 |
| KIRC | kidney | 0.916 | 0.574 | 0.627 |
| KIRP | kidney | 1.07 | 0.51 | 0.475 |
| LIHC | liver | 0.585 | 0.37 | 0.632 |
| LUAD | lung | 1 | 0.168 | 0.168 |
| LUSC | lung | 1.03 | 0.444 | 0.431 |
| OV | ovary | 1.81 | 0.584 | 0.322 |
| PAAD | pancreas | 2.18 | 2.39 | 1.1 |
| PRAD | prostate | 0.964 | 0.313 | 0.324 |
| READ | rectum | 0.829 | 0.311 | 0.375 |
| SKCM | skin | 0.497 | 1.04 | 2.1 |
| STAD | stomach | 0.818 | 0.435 | 0.532 |
| TGCT | testis | 1.81 | 0.857 | 0.474 |
| THCA | thyroid gland | 1.14 | 0.216 | 0.189 |
| UCEC | endometrium | 1.47 | 0.245 | 0.167 |

**Figure 2:**
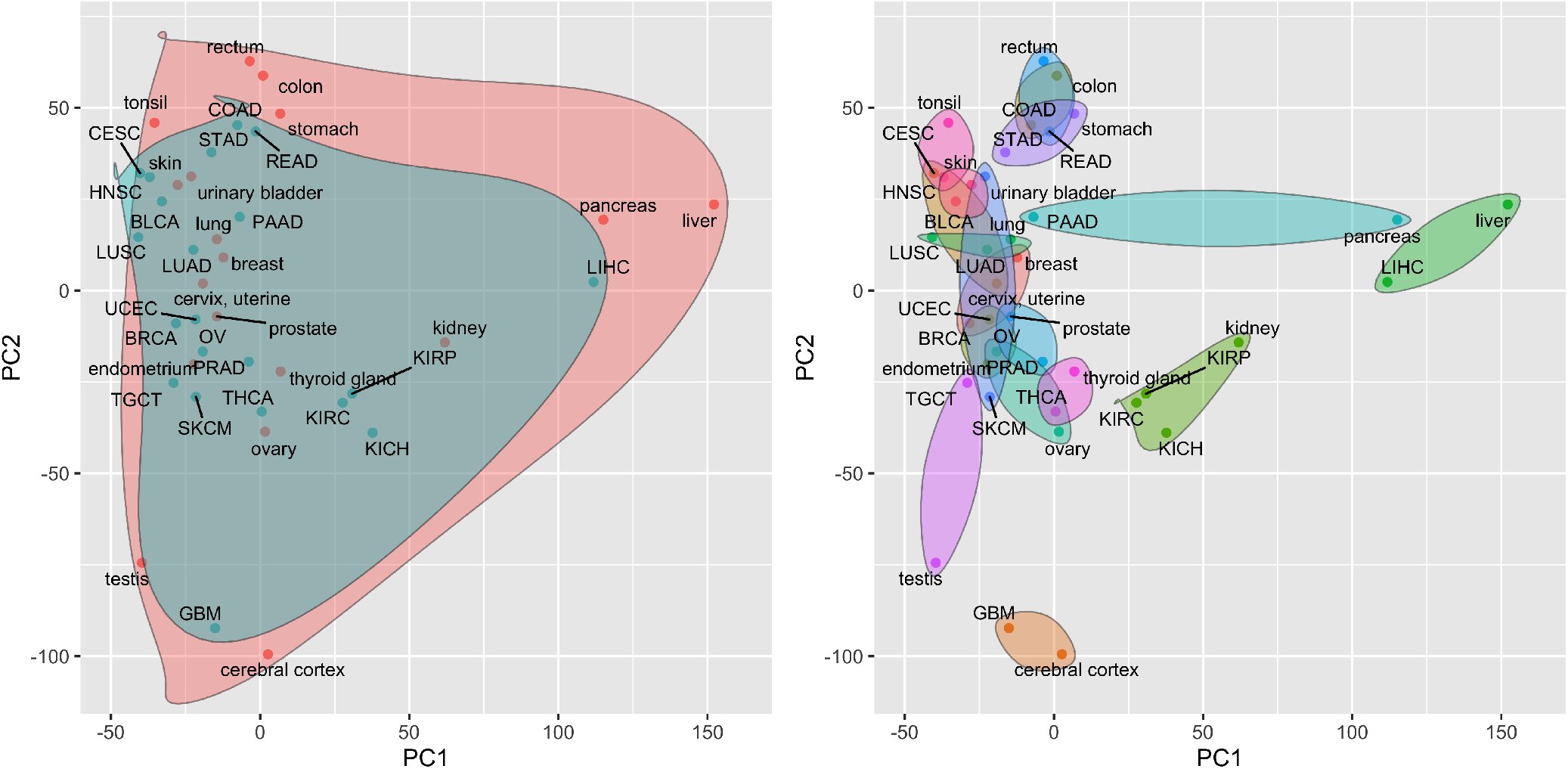
Projection of TCGA cancer types and associated HPA normal tissues onto the first two principal components. Principal components are computed from a matrix of tissue-specific and cancer-specific mean gene expression values, i.e., a matrix containing all 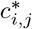 and 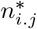 statistics. a) Cancers are represented by blue points and are enclosed in the blue shaded region; normal tissues are represented by red points and are enclosed in the red shaded region. b) Each normal tissue and the associated cancer type(s) are enclosed in a separate shaded region.

### 2.4 Association between normal tissue-specificity, cancer/normal differential expression and cancer survival

The ratio of cancer to normal tissue gene expression is inversely associated with normal tissue specificity for most of human genes in all tested cancer/normal pairs. In other words, tissue-specific genes tend to be down-regulated in cancer, which is consistent with a view of cancer as a dedifferentiation process. This finding is quantified by the *ρ_case/ctrl_* values in Table 4, which capture the rank correlation between normal tissue specificity 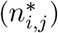 and cancer/normal relative expression (*r_i,j_*) and are negative for all 21 profiled cancer types. A related finding is that normal tissue specificity tends to be positively associated with favorable survival in cancer, i.e., an increase in expression of normal tissue specific genes is associated with improved survival in cancer. Genes that are not tissue-specific have the inverse association, i.e., an increase in expression is associated with worse survival. Genes that are down-regulated in a tissue relative to other tissues tend to have no survival association. The rationale behind these survival associations is that cancer cells are more dedifferentiated than normal tissue cells [8]. This survival association are quantified by the *ρ_surv_* values in Table 4, which capture the rank correlation between normal tissue specificity 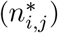 and prognostic ability (*s_i,j_*) and are near zero or positive for all 21 profiled cancer types. Figure 3 provides a more detailed visualization of the association of normal tissue-specificity with cancer/normal relative expression and cancer survival for liver cancer.

**Table 4:**
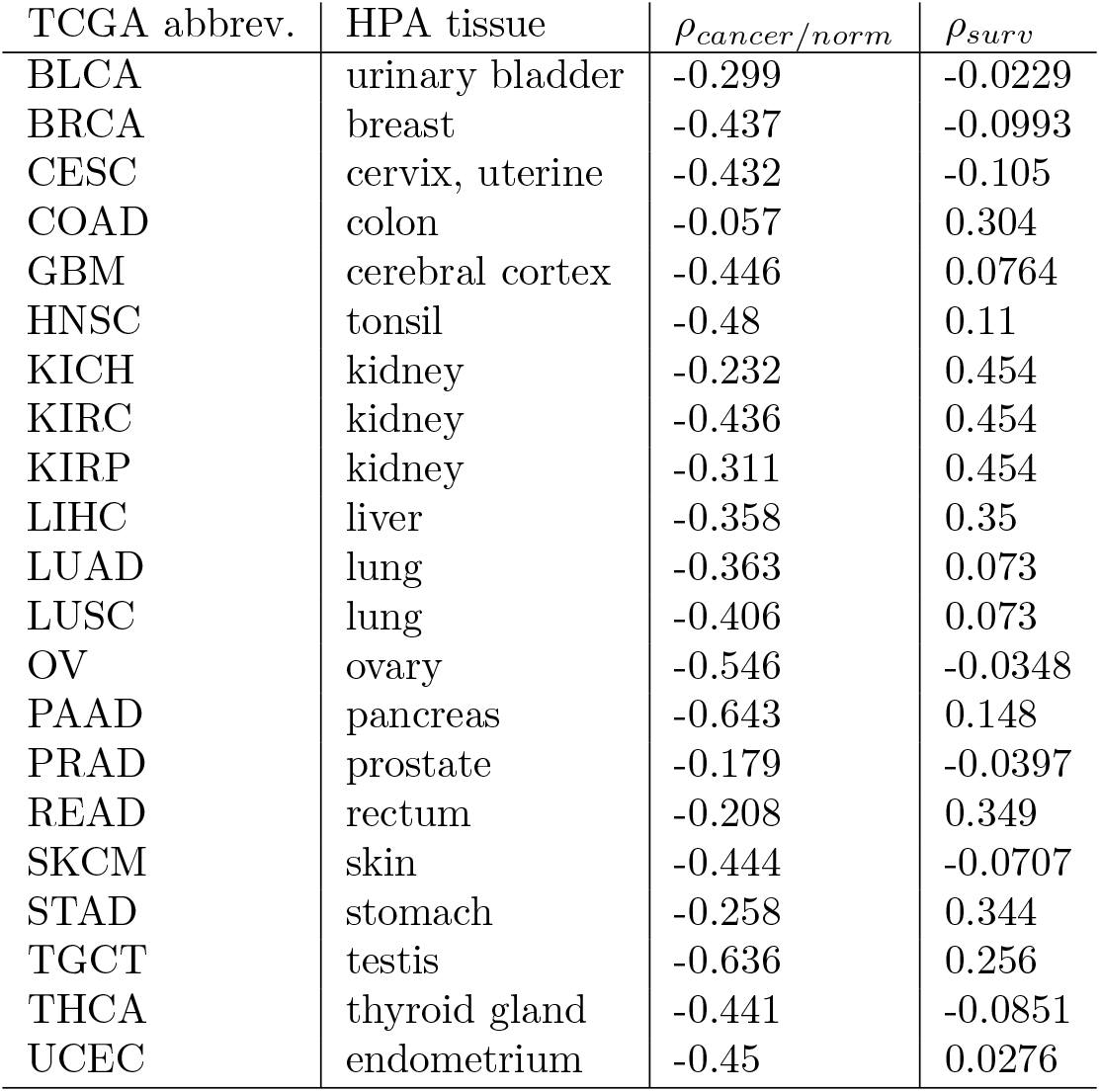
Correlations between normal tissue-specific gene weights and either cancer/normal relative expression or an indicator of cancer prognostic ability. The *ρ_cancer/norm_* column holds the Spearman rank correlation between normal tissue-specific gene weights 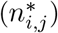 and the log ratio of mean expression in the cancer type to mean expression in the normal tissue (*r_i,j_*). The *ρ_surv_* column holds the Spearman rank correlation between normal tissue-specific gene weights 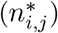 and the signed log of the p-value from a Kaplan-Meir test of the association between gene expression and cancer survival as computed by Uhlen et al. [8]) (*s_i,j_*, which is computed as −log(p-value) for favorable genes and log(p-value) for unfavorable genes).

**Figure 3:**
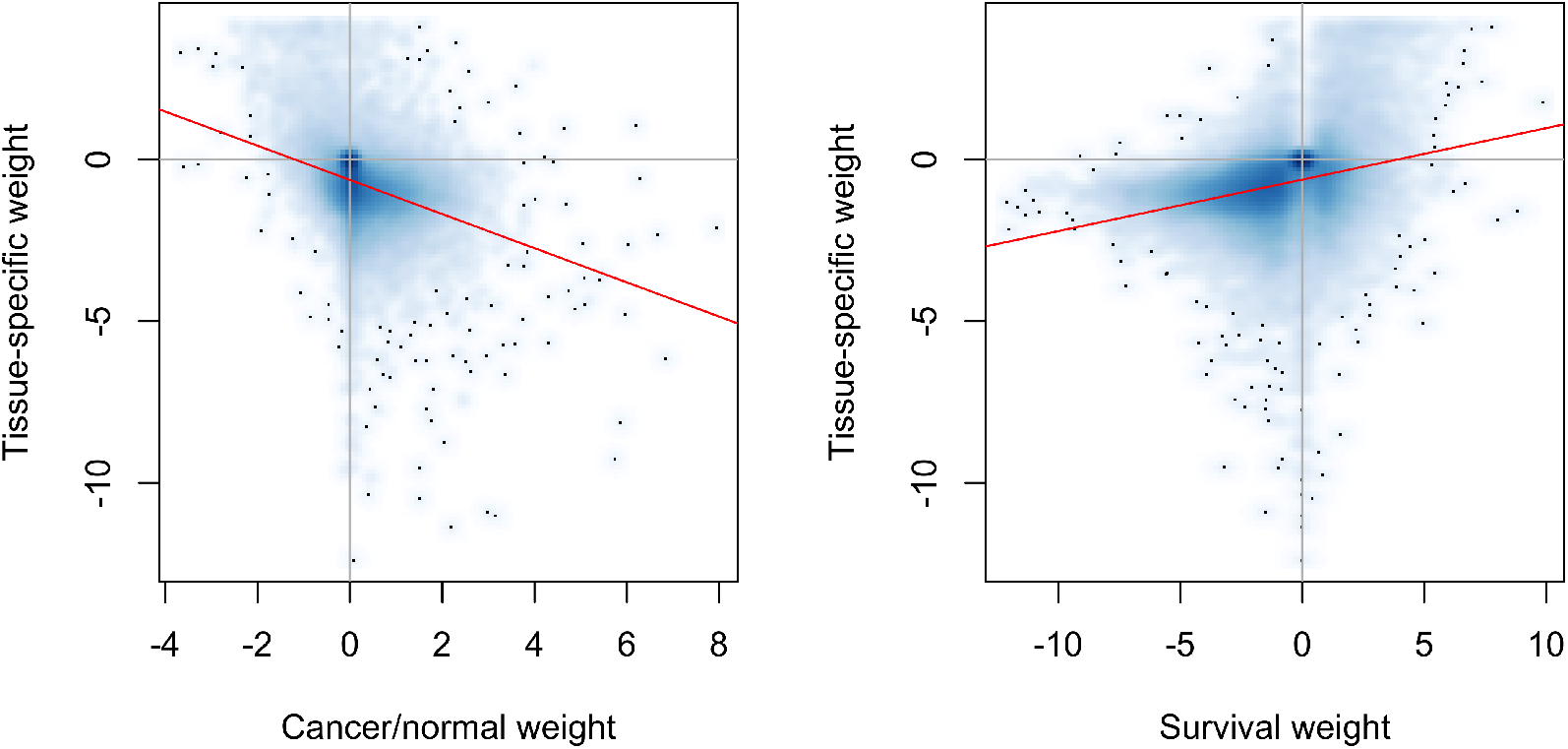
Association of liver-specificit 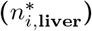 liver cancer/normal liver differential expression (*r_i,_*_LIHC_) and liver cancer prognostic value (*s_i,_*_LIHC_). Each point in both plots represents a single gene and the red lines reflect the linear regression fit. a) Association between liver-specificity and differential expression between liver cancer and normal liver. b) Association between liver-specificity and prognostic value of gene expression for liver cancer.

### 2.5 Using normal tissue gene activity to improve the comparative analysis of cancers

As illustrated in Figure 2 and Table 3, the relative expression of genes between different cancer types is associated with the relative expression in the corresponding normal tissues. This association can be leveraged to improve the comparative analysis of cancer types based on transcriptomic data. Specifically, when attempting to identify genes that are differentially expressed (DE) between two cancer types, genes that are DE between the associated normal tissues can be prioritized, e.g., via gene filtering or a hypothesis weighting scheme like weighted false discovery rate (wFDR) [11]. Although this will improve the power for detecting DE genes between cancer types, the results will highlight differences between the underlying normal tissues rather than differences between the cancer types that are independent of normal tissue physiology. Alternatively, removing tissue-specific genes (or gene sets) can enable the identification of differences between cancer types that are due to malignant processes and not differential gene activity within the normal tissues. This second approach is one we have successfully used in the past for the analysis of gene expression in primary colorectal tumors and colorectal metastatic lesions in the liver and lung [12]. The pan cancer impact of this approach is illustrated in Figure 4 and Table 5. Figure 4 visualizes the rank correlation between the fold-change in gene expression between each pair of cancers and the corresponding pair of normal tissues. For cells below the diagonal, all genes are used to compute the correlations; for cells above the diagonal, genes that have a log2 fold-change between normal tissues of less than 0.5 or more than 2 are excluded. The positive correlation values generated on all genes indicate that the pattern of DE between most cancers mirrors the pattern of DE found between the corresponding normal tissues. The lower correlation values in the cells above the diagonal confirms that filtering out genes showing large DE between associated normal tissues will generate cancer DE results that are distinct from the normal tissue biology. Table 5 demonstrates the impact of this association for the comparative analysis of liver cancer and glioblastoma. This table lists the 10 Molecular Signatures Database (MSigDB) [13] Hallmark pathways most significantly associated with relative gene expression between liver and cerebral cortex and between liver cancer and glioblastoma. Gene set testing of liver cancer/glioblastoma relative expression was performed using all genes (middle column) and using genes filtered according to DE between normal liver and cerebral cortex (right column). Filtering for this analysis removed all genes whose log2 fold-change in expression between liver and cerebral cortex was *≤ −*2 or *≥* 2. As seen in the table, differential pathway activity between liver cancer and glioblastoma is similar to differential pathway activity between normal liver and cerebral cortex when all genes are considered. When genes exhibiting significant normal tissue DE are removed, the comparative cancer results are distinct from the normal tissue results.

**Table 5:**
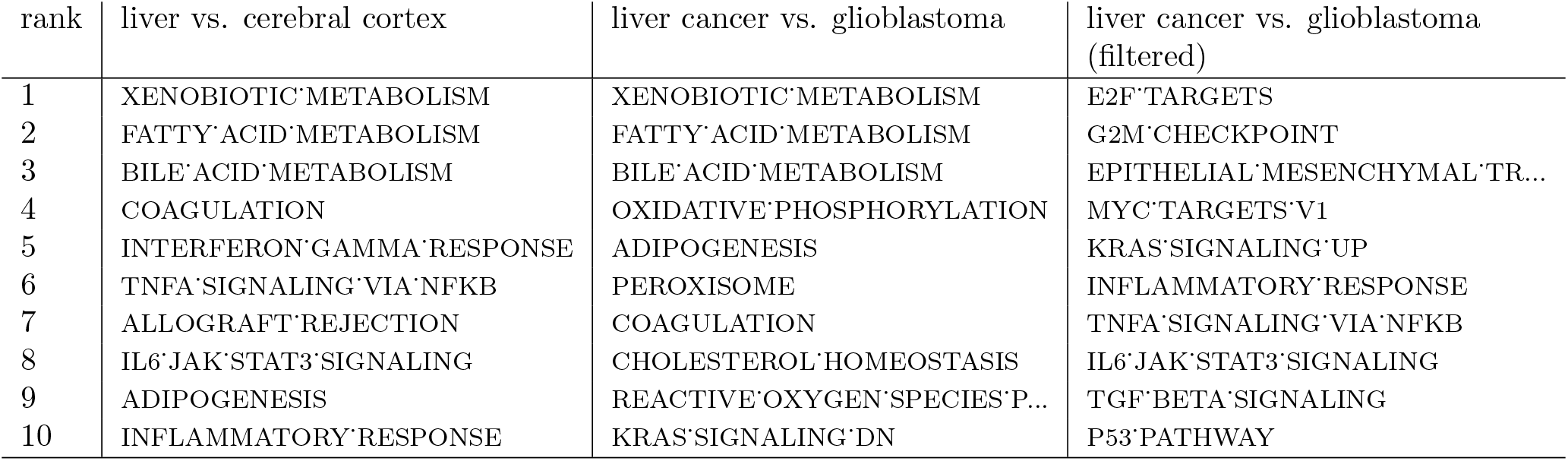
Impact of normal tissue-specificity on comparative analysis of liver cancer and glioblastoma. The table lists the 10 MSigDB Hallmark pathways most significantly associated with relative gene expression between liver and cerebral cortex and between liver cancer and glioblastoma. Gene set testing of cancer relative expression was performed using all genes (middle column) and using genes filtered according to normal tissue differential expression (right column). Filtering specifically removed all genes whose log2 fold-change in expression between liver and cerebral cortex was ≤ −2 or ≥ 2. Without filtering, differential gene expression between liver cancer and glioblastoma is similar to differential expression between normal liver and cerebral cortex.

| rank | liver vs. cerebral cortex | liver cancer vs. glioblastoma | liver cancer vs. glioblastoma (filtered) |
| --- | --- | --- | --- |
| 1 | XENOBIOTIC'METABOLISM | XENOBIOTIC'METABOLISM | E2F'TARGETS |
| 2 | FATTY'ACID'METABOLISM | FATTY'ACID'METABOLISM | G2M'CHECKPOINT |
| 3 | BILE'ACID'METABOLISM | BILE'ACID'METABOLISM | EPITHELIAL'MESENCHYMAL'TR... |
| 4 | COAGULATION | OXIDATIVE'PHOSPHORYLATION | MYC'TARGETS'V1 |
| 5 | INTERFERON'GAMMA'RESPONSE | ADIPOGENESIS | KRAS'SIGNALING'UP |
| 6 | TNFA'SIGNALING'VIA'NFKB | PEROXISOME | INFLAMMATORY'RESPONSE |
| 7 | ALLOGRAFT'REJECTION | COAGULATION | TNFA'SIGNALING'VIA'NFKB |
| 8 | IL6'JAK'STAT3'SIGNALING | CHOLESTEROL'HOMEOSTASIS | IL6'JAK'STAT3'SIGNALING |
| 9 | ADIPOGENESIS | REACTIVE'OXYGEN'SPECIES'P... | TGF'BETA'SIGNALING |
| 10 | INFLAMMATORY'RESPONSE | KRAS'SIGNALING'DN | P53'PATHWAY |

**Figure 4:**
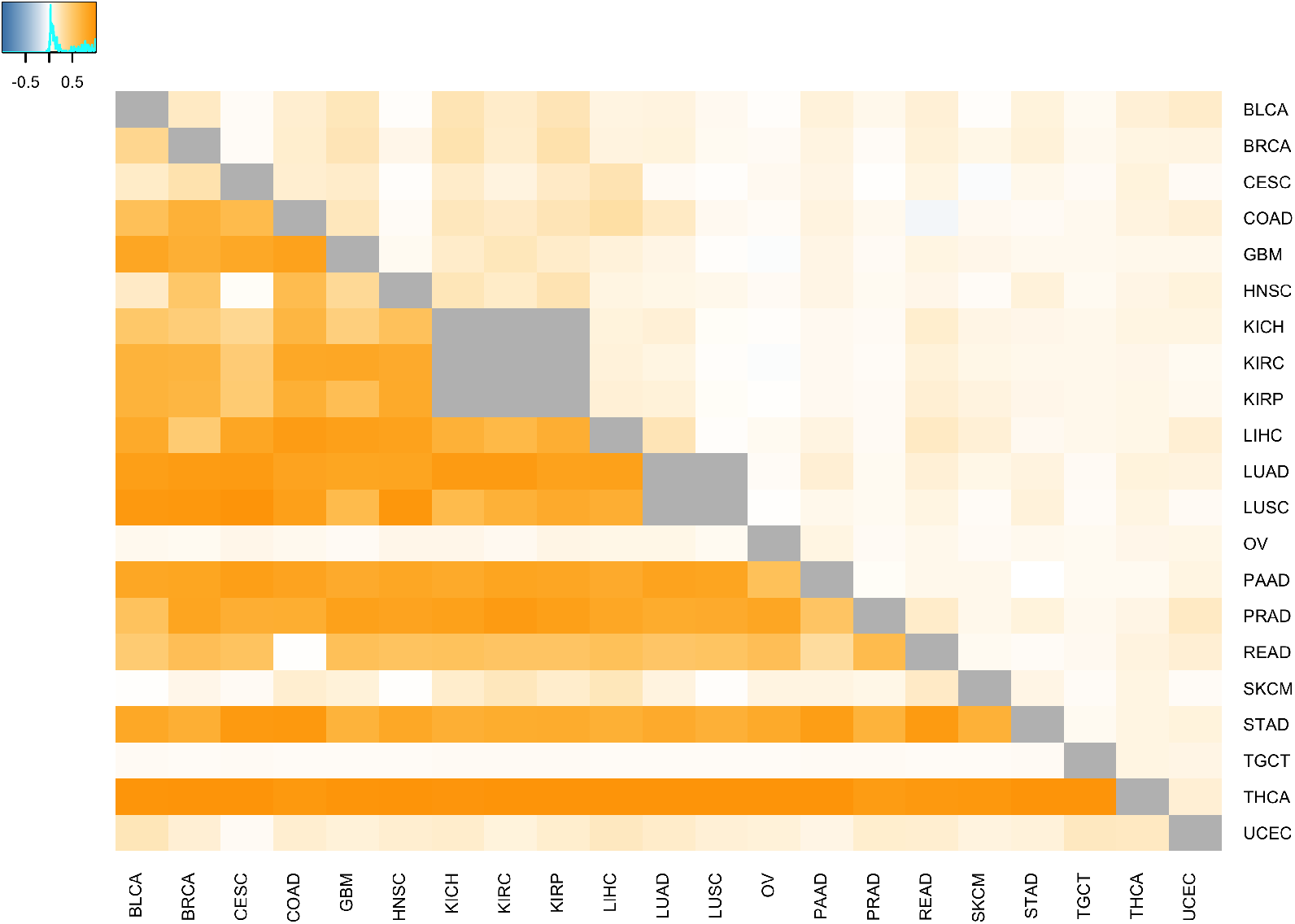
Impact of normal tissue-specificity on cancer differential expression. Cell coloring reflects the Spearman rank correlation between the fold-change in gene expression between each pair of cancers and the corresponding pair of normal tissues. Large correlations indicate that the expression difference between cancers mirrors the expression difference in the normal tissues. For cells below the diagonal, all genes are used to compute the correlations; for cells above the diagonal, genes that have a log2(fold-change) between normal tissues of less than 0.5 or more than 2 are excluded. As indicated by the lower correlations in cells above the diagonal, filtering out genes that show large differential expression between normal tissues will generate cancer differential expression results that are distinct from the normal tissue biology.

### 2.6 Use of normal tissue-specificity to improve the comparative analysis of normal and neoplastic tissue

As illustrated in Table 4, tissue-specific genes are more likely to be down-regulated in cancer as compared to the associated normal tissue and non-specific genes are more likely to be up-regulated; these associations hold across all 21 evaluated cancer types. Similar to the approach outlined in Section 2.5 for the comparative analysis of different cancers, this association between normal tissue-specificity and cancer/normal DE can be used to prioritize genes for analysis via either gene filtering or hypothesis weighting. Figure 5 illustrates the increase in power that can be achieved by filtering genes according to normal tissue-specificity prior to a cancer/normal DE analysis. Specifically, this figure contains quantile-quantile (Q-Q) plots of p-values from gene set testing of MSigDB Hallmark pathways relative to cancer/normal relative expression (i.e, the *r_i,j_* statistics) for the 21 profiled cancer types using all genes or genes filtered to remove the 20% of genes with the most extreme tissue-specificity values (i.e., the 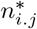 statistics). To test for pathways enriched in genes that are up-regulated in the cancer as compared to the associated normal tissue (panel a), filtering was performed to remove the most tissue-specific genes. To test for pathways enriched in genes that are down-regulated in the cancer as compared to the associated normal tissue (panel b), filtering was performed to retain tissue-specific genes.

**Figure 5:**
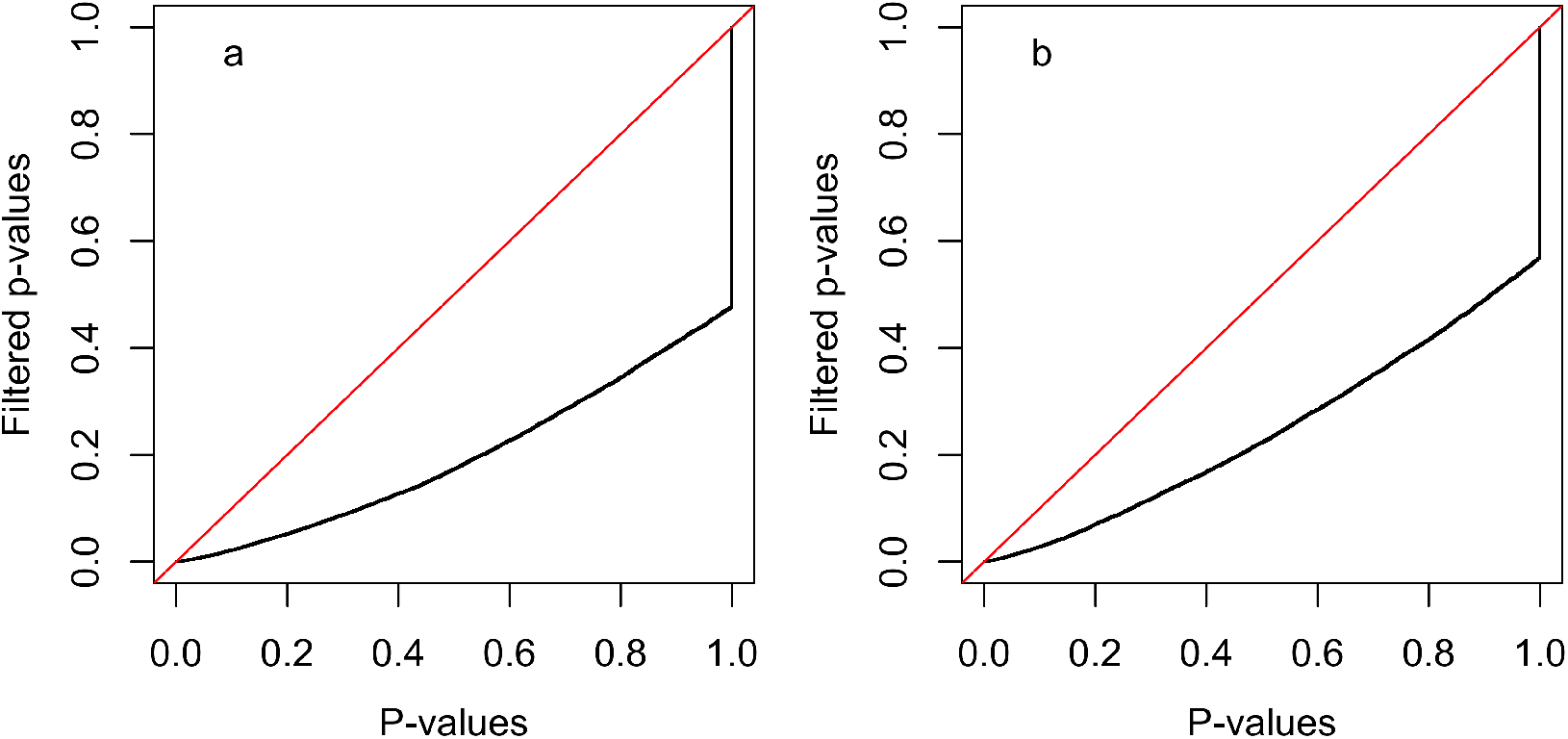
Quantile-quantile plots illustrating the impact of tissue-specific gene filtering on gene set testing of MSigDB Hallmark pathways using cancer/normal relative expression (*r_i,j_*). Each plot contrasts the distribution of p-values (the black line) from tests for all 50 Hallmark pathways for each of the 21 cancer/normal pairs (total of 1050 tests) using all genes or genes filtered according to tissue-specificity. The red line has a slope of 1.0 and captures the distribution that would be expected if filtered and un-filtered p-values had the same distribution. In both panels, the x-axis reflects the p-value distribution when all genes are included and the y-axis reflects the p-value distribution when genes are filtered according to normal tissue-specificity 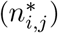. Panel a) contains the results from tests for enrichment of large *r_i,j_* statistics among the gene set members (i.e., are gene set members more likely to be up-regulated in cancer vs. the normal tissue?) and gene filtering removes the 20% of genes with the largest tissue-specificity values (i.e., keeps genes that are less tissue-specific). Panel b) contains the results from tests for enrichment of small *r_i,j_* statistics among the gene set members (i.e., are gene set members more likely to be down-regulated in the cancer vs. the normal tissue?) and gene filtering removes the 20% of genes with the smallest tissue-specificity values (i.e., keeps genes that are more tissue-specific). For both panels, the p-values for gene sets whose direction of enrichment is opposite the target direction are set to 1.0, which causes the vertical portion of the black line. As illustrated in the Q-Q plots, gene filtering in both cases improves gene set testing statistical power.

### 2.7 Use of normal tissue-specificity to improve cancer survival analysis

Normal tissue-specificity can also be used to improve the power of cancer survival analysis. As shown in Table 4, tissue-specific genes are more likely to be favorably prognostic for cancer survival with non-tissue-specific genes more likely to be unfavorably prognostic. Unlike the association between tissue specificity and cancer/normal DE, the association with survival is only pronounced in a subset of the profiled cancer types. For these cancer types, an approach similar to those outlined in Sections 2.5 and 2.6 can be used to prioritize genes for survival analysis. Figure 6 illustrates the increase in survival analysis power that can be achieved by filtering genes according to normal tissue-specificity. Similar to Figure 5, this figure contains quantile-quantile (Q-Q) plots of p-values from survival analyses for all 21 profiled cancer types using all genes or a subset of genes filtered according to normal tissue-specificity (the 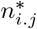 statistics). In this case, the p-values are generated via Kaplan-Meir (KM) tests of the association of gene expression with cancer survival following the approach of Ulhen et al. [8] (i.e., the *s_i,j_* statistics). To identify favorably prognostic genes (panel a), filtering was performed to remove genes where 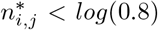, i.e., genes down-regulated in the associated tissue. To identify unfavorably prognostic genes (panel b), filtering was performed to remove genes where 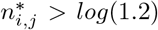,i.e., genes up-regulated in the associated tissue.

**Figure 6:**
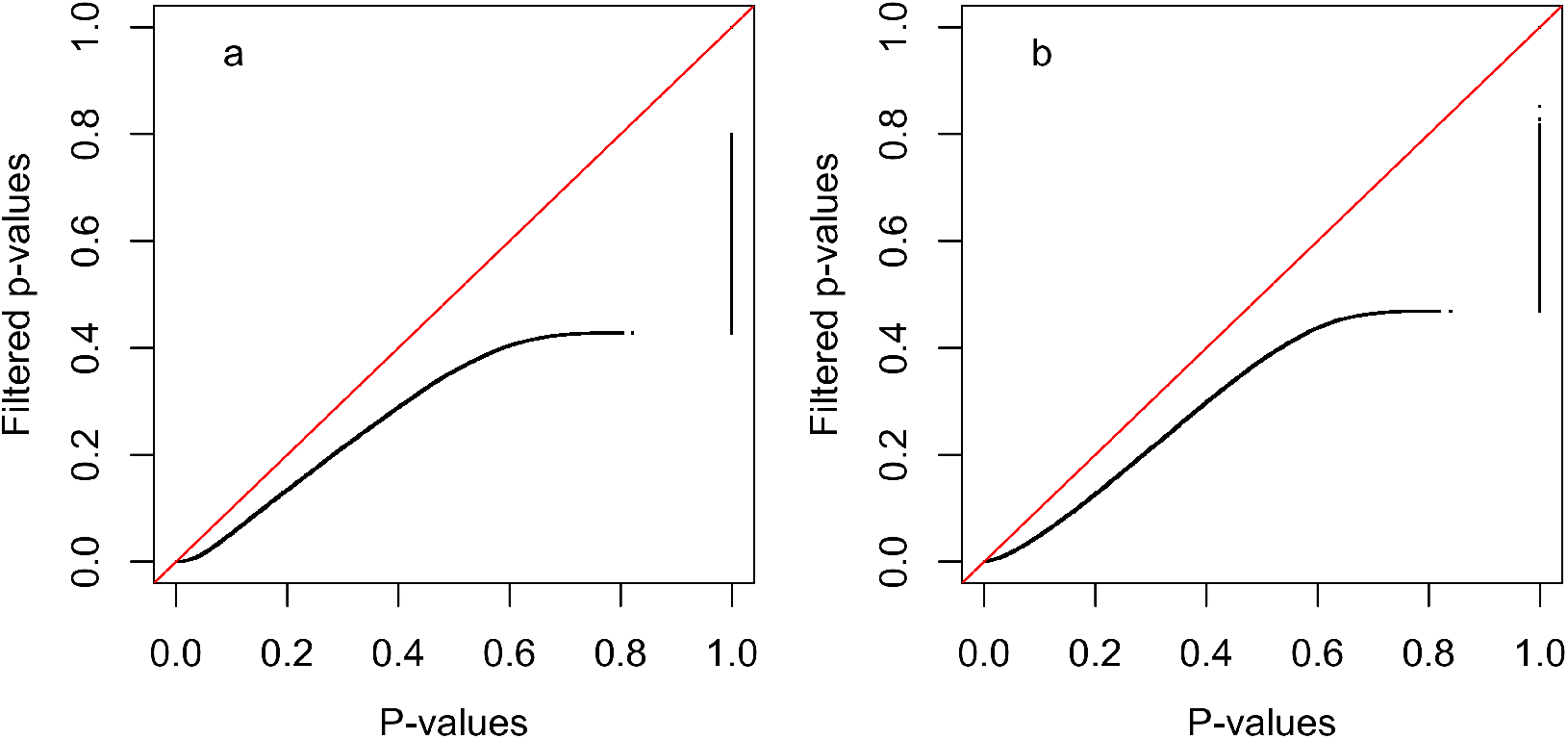
Quantile-quantile plots illustrating impact of tissue-specific gene filtering on the distribution of p-values from Kaplan-Meir (KM) tests of the association of gene expression with cancer survival as computed using the approach of Ulhen et al. [8] (the *s_i,j_* statistics). Each plot contrasts the distribution of p-values (the black line) for all genes for each of the 21 cancer types or for genes filtered according to tissue-specificity. The red line has a slope of 1.0 and captures the distribution that would be expected if filtered and un-filtered p-values had the same distribution. In both panels, the x-axis reflects the p-value distribution when all genes are included and the y-axis reflects the p-value distribution when genes are filtered according to normal tissue-specificity 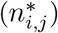. Panel a) contains the results when only favorably prognostic genes are considered (i.e., genes where increased expression is associated with better survival; p-values for unfavorable genes are set to 1.0) and gene filtering removes genes where 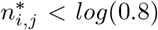 (i.e., removes genes expressed at a lower level in the associated normal tissue than in the average tissue). Panel b) contains the results when only unfavorably prognostic genes are considered (i.e., genes where increased expression is associated with worse survival; p-values for favorable genes are set to 1.0) and gene filtering removes genes where 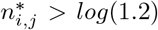(i.e., removes genes expressed at a higher level in the associated normal tissue than in the average tissue). As illustrated in the Q-Q plots, gene filtering in both cases improves survival analysis statistical power.

## 3 Conclusions

The biology of human cancer is highly tissue-specific with the majority of cancer-related somatic alterations occurring in only a small number of tissue types and the functional impact of common and inherited mutations frequently exhibiting a tissue-specific functional impact. An important consequence of cancer tissue-specificity is that the biology of normal tissues holds important information regarding the molecular features of associated cancers, information that can be leveraged to improve the power and accuracy of cancer genomic analyses. To date, most research exploring the joint analysis of normal tissue and cancer genomic data has focused on the analysis of tumor and adjacent normal samples. The development of approaches that leverage the general characteristics of normal tissues for cancer analysis has only received limited attention with most investigations focusing on specific alterations within a single cancer type. To address this research gap and support use cases where adjacent normal tissue samples are unavailable, we have explored the genome-wide association between the transcriptomes of 21 solid human cancers profiles by TCGA and their associated normal tissues as profiled in healthy individuals by the HPA. Although the mean transcriptomic profiles of normal and cancerous tissue appear distinct, a strong association is revealed between each cancer and the corresponding normal tissue when gene expression data is transformed into tissue or cancer-specific values, i.e., the ratio of expression in one tissue or cancer relative to the mean in other tissues or cancers. As we have demonstrated through the analysis results presented in this paper, the strong association between cancer-specific and tissue-specific gene expression can be leveraged to significant improve statistical power and biological interpretation for cancer survival analysis, cancer comparative analysis, and analysis of cancer/normal pairs.

